# Mitochondrial dynamics can resolve stoichiometric challenges in plant mtDNA repair

**DOI:** 10.64898/2026.10.08.757513

**Authors:** Iain G. Johnston, Daniel B. Sloan, Ian Max Møller

## Abstract

Although mitochondrial DNA (mtDNA) is subject to damage and high mutation rates in many eukaryotes, plants have a variety of mtDNA repair pathways that somehow result in exceptionally low mutation rates. However, the stoichiometry of biomolecules in plant mitochondria raises challenges for the DNA repair process. The protein complexes responsible for repair pathways contain subunits present at low copy numbers per mitochondrion. Mitochondria on average contain less than a full genome, limiting the capacity for templated repair, and raising the risk that repair machinery is localised to a DNA-empty mitochondrion where it cannot be used. Here we build on recent experimental characterisation of the copy numbers and physical processes involved to model the capacity for DNA repair in plant mitochondria. We find that targeted import of protein machinery, biased fusion of mitochondria bearing DNA, and exchange of biomolecules upon fusion can improve repair capacity through different mechanisms. Specifically, our model indicates that base excision repair can keep ∼95% of mtDNA intact even without targetting or exchange rules and that targetting protein import to DNA-bearing mitochondria or exchanging protein content drives this to effectively 100%. In contrast, template repair cannot keep a majority of mtDNA intact unless mtDNA is shared across mitochondria through fusion, in which case its efficacy increases. Exchanging of metabolons (repair protein complexes) increases repair efficiency across all circumstances. We discuss the implications of our model for plant development and organelle inheritance.

## Introduction

Plant mitochondria are essential for the respiratory and bioenergetic processes that feed the world (Chustecki & Johnston, 2024). As in other eukaryotes, plant mitochondrial DNA (mtDNA) encodes vital bioenergetic machinery that contributes to the proper functioning of mitochondria. This mtDNA can become damaged, compromising cellular function. Because of the centrality of mitochondrial function to plant (and eukaryotic) life, a collection of repair and maintenance mechanisms exist to mitigate mtDNA damage (Chevigny et al., 2020; Gualberto & Newton, 2017; Jeppesen et al., 2011; Schatz-Daas et al., 2022; Veeraragavan et al., 2024).

Plant mitochondria have several characteristics that complicate these mechanisms. Unlike animal and yeast mitochondria, plant mitochondria rarely undergo large-scale fusion, with exceptions in the shoot apical meristem and the early stages of germination (Paszkiewicz et al., 2017; Segui-Simarro et al., 2008). In most tissues, fragmented mitochondria undergo many transient fusion-fission events acting to share content through the mitochondrial population (Arimura et al., 2004; Chustecki et al., 2021; Giannakis et al., 2022; Logan, 2010). This fusion-fission is facilitated by collective mitochondrial movement, by which mitochondria meet many potential exchange partners as they travel around the cell (Arimura, 2018; Arimura et al., 2004; Chustecki et al., 2021, 2022), leading to mixing throughout the population on the timescale of hours (Arimura et al., 2004). Many plant mitochondria within vegetative tissues lack mtDNA entirely, with fewer mtDNAs than mitochondria in plant cells (Chustecki et al., 2025; Preuten et al., 2010); mitochondria without mtDNA appear to move less and undergo fewer exchanges (Chustecki et al., 2025).

On the background of this physical system, the genetic population of mtDNA in the plant cell is shaped by several different processes. Base excision repair (BER) (Ferrando et al., 2019; Jeppesen et al., 2011) acts in plants (Boesch et al., 2009) along with several other pathways to repair damage to individual mtDNA molecules; template repair (TR) (Broz et al., 2022, 2024; Edwards et al., 2021; Peñafiel-Ayala et al., 2026; Wu et al., 2020; Zhou et al., 2026; Zwonitzer et al., 2024), a recombination-mediated process, acts to repair damage on one mtDNA by using another as a template. Both these processes involve collections of steps, each catalysed by a protein. The plant chondriome, comprising all the mitochondria in a cell, contains 2000-2500 different proteins depending on the cell type and the environmental conditions (Jiang et al., 2021; Kim et al., 2026; Rao et al., 2017). Estimates from dark-cultured Arabidopsis cell suspensions indicate that the relative abundance of proteins varies over at least five orders of magnitude, from the most abundant voltage-dependent anion channel 1 with 44 381 copies per medium-sized mitochondrion with around 1.4 million protein molecules to less than one copy per mitochondrion (50 different proteins) (Fuchs et al., 2020).

The average per-mitochondrion stoichiometry of proteins involved in mtDNA repair is surprisingly low in several important cases. Many of the proteins are part of protein complexes that only function if all the components are present (Rugen et al., 2021; Senkler et al., 2017). DNA-binding proteins generally have low copy numbers in plant mitochondria (Fuchs et al., 2020), and since mtDNA is strongly substoichiometric (Preuten et al., 2010), the co-localization of the DNA-binding proteins with the mtDNA may be rare, if it occurs randomly. For example, the TR pathway requires the simultaneous presence of two mtDNA molecules, the damaged mtDNA and an intact template mtDNA, as well a collection of proteins, some of which are present only at very low average copy numbers per mitochondrion. A specific protein example is MSH1, the first protein/enzyme in the TR pathway, which is present on average as a single dimer per organelle (Fuchs et al., 2020)).

Given that mitochondria may only contain limited numbers of mtDNA and repair proteins, it is far from guaranteed (under a random arrangement of biomolecules) that every mtDNA is capable of being repaired. Indeed, it has been suggested that organelle DNA is effectively abandoned in some vegetative plant tissues (Oldenburg & Bendich, 2015). Here, we combine mathematical and simulation modelling with recent data characterising per-mitochondrion stoichiometry to demonstrate conditions and mechanisms that could support the repair and maintenance of plant mtDNA despite these limited copy numbers.

## Methods

Broadly, our model consists of a collection of mitochondria, which can contain different numbers of different proteins (which turn over with time) and mtDNA (which becomes randomly damaged over time). MtDNA damage can be repaired by the action of proteins. Different targetting, fusion, and exchange rules, respectively, determine how proteins are imported into mitochondria, which mitochondria undergo fusion, and which biomolecules are exchanged upon fusion-fission events.

### Model basics

A cell in our model (Fig. 1A) is pictured as containing *n_M_* mitochondria. We consider a protein complex (which will act as a metabolon) that consists of *n* subunit proteins. The copy number of each of these subunits, and mtDNA molecules, in mitochondrion *i* is described by a vector *c*^(*i*)^ (where elements 1 through *n* describe copy numbers of each protein) and a scalar describing the mitochondrion’s copy number of mtDNA. Initially, *n_M_*/3 mtDNA molecules (Fuchs et al., 2020; Preuten et al., 2010) are randomly assigned across the mitochondria in the cell. mtDNA may be intact or damaged. Intact mtDNA becomes damaged randomly with rate *μ*. Once damaged, mtDNA can be returned to the intact state through the action of repair protein complexes.

**Figure 1.**
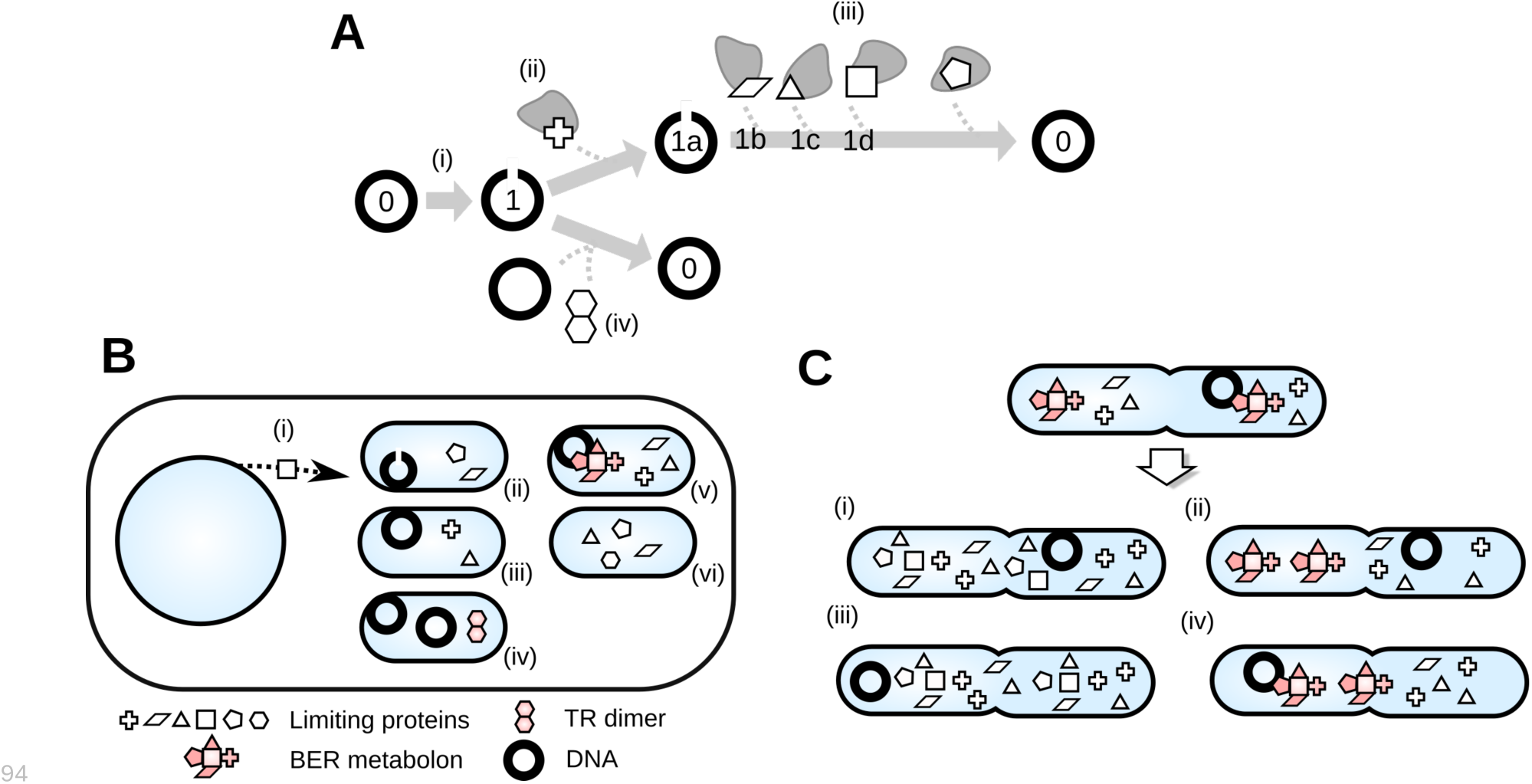
Summary of model behaviour. **(A)** Damage and repair to mtDNA. (i) Intact mtDNA (labelled 0) can become damaged (labelled 1) with some damage rate. (ii) The BER-like pathway starts with recognition of damage by a protein subunit, leading to state 1a. (iii) Subsequent subunits in the repair pathway drive the repair process through different states 1b, 1c, 1d, and the final step produces an intact mtDNA. (iv) The TR-like pathway involves recognition and repair using a protein dimer and an additional copy of mtDNA. **(B)** Compartments, molecules, and import. We consider only the “limiting” subunits for each complex – those present at lowest per-mitochondrion copy number. Other subunits are assumed to be present in excess. (i) Subunits are produced by the nucleus and imported into mitochondria either randomly or according to a targetting rule. This can be (ii) *Damaged* (mitochondria containing damaged mtDNA); (ii)-(v) *DNA-bearing* (mitochondria containing any mtDNA); (ii),(iii) *DNA-bearing, incomplete* (mitochondria with mtDNA but without a full set of repair proteins); (ii), (iii), (vi) *Incomplete* (mitochondria without mtDNA and/or a full repair set). **(C)** Exchange on fusion. When two mitochondria fuse and fission, different exchange rules determine which biomolecules are “pooled” and shared between the resultant daughters. Example rules: (i) *Subunits* (all protein subunits are mixed), (ii) *Metabolons* (complete sets of repair proteins, and remainder subunits, are mixed); (iii) *DNA + Subunits* (individual mtDNA molecules and protein subunits are mixed); (iv) *DNA-metabolons + Metabolons* (DNA-metabolon complexes and metabolons without mtDNA are mixed).

### Repair protein dynamics

In our model, repair protein complexes can convert damaged mtDNA to intact mtDNA through two pathways. The first, BER-like pathway, involves a series of steps, each accomplished by a different protein subunit acting on a single damaged mtDNA. The second, TR-like pathway, involves one rate-limiting step, accomplished by a protein dimer acting on a damaged mtDNA in the same compartment as another mtDNA (Fig. 1). This model is based on a putative MSH1 pathway that involves introduction of a double strand break (DSB) at damaged sites, followed by recombination repair of the DSB (Broz et al., 2022; Christensen, 2014; Zhou et al., 2026). Because the proteins involved in subsequent recombinational steps appear to be present at much greater copy numbers (including members of the OSB, REC, WHY families (Fuchs et al., 2020)), we do not consider the rate of these steps to be limited by protein stoichiometry. We also note that it is unclear if and how MSH1 interacts with single mtDNA molecules in the absence of an mtDNA partner, at it could potentially induce DSBs without an available template for immediate repair. For the purposes of this model, we ignore any potential DSB-inducing activity when only a single mtDNA molecule is present, but this represents an important area of uncertainty in the current literature.

Schematically, these processes resemble (Fig. 1A):

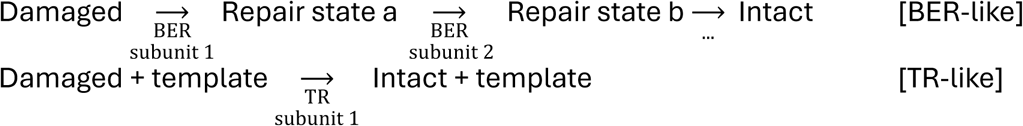

Although each step is generally catalysed by a subunit protein, we consider only the lowest-copy-number component of each complex as the stoichiometric limiting factor in these processes, assuming that other subunits are present in excess.

The cell produces proteins at rate *λ*. Each protein is imported by a mitochondrion with rate *ρ* according to a *targetting rule* (Fig. 1Ai). These targetting rules are: *Damaged* (only target to mitochondria containing damaged mtDNA); *DNA-bearing* (only target to mitochondria containing mtDNA); *DNA-bearing, incomplete* (only target to mitochondria with mtDNA but without a full set of repair proteins); *Incomplete* (only target to mitochondria without a combination of mtDNA and a full set of repair complexes); or *Random* (target each protein to a random mitochondrion). Proteins turn over with rate *ν*, reflecting their damage, removal, and replacement.

### Mitochondrial fusion and exchange dynamics

Mitochondrial fusion events occur with rate *κ* per cell per unit time. Two rules determine behaviour upon fusion. The *social rule* determines which mitochondria fuse – this is either *All mitochondria*, or *DNA-bearing* (only mitochondria contain mtDNA fuse). The *exchange rule* (Fig. 1C) determines which biomolecules are mixed upon fusion. This can be *Metabolons* (complete sets of repair proteins are mixed), *DNA + Subunits* (individual mtDNA molecules and protein subunits are mixed); *DNA-metabolons* (complexes of mtDNA and complete sets of repair proteins are mixed); *DNA-metabolons + Metabolons* (DNA-metabolon complexes and metabolons without mtDNA are mixed); *None* (no mixing); *Subunits* (individual protein subunits are mixed). After mixing, fused mitochondria immediately fission, modelling “kiss-and-run” behaviour (Arimura, 2018; Logan, 2010).

### Parameter values

Experimental evidence guides choices of some parameters. We take *n_M_* = 300, broadly matching observations from microscopy (Chustecki et al., 2021, 2025; Logan, 2010). Mixing of mitochondrial matrix content through the cellular population (through fusion) occurs on the timescale of 1-2 h in onion epidermis (Arimura et al., 2004). This corresponds to every mitochondrion having undergone at least one fusion event. The average number of fusion events per mitochondrion is 2 *κ t* / *n_M_*, so *κ* = *n_M_*/2 = 150 per unit time achieves this mixing rate. Across a collection of functions and families, plant proteins have a mean turnover rate around 0.1 per day (Nelson et al., 2013), so we use this turnover rate for our model proteins.

The DNA damage rates that appear in the scientific literature are the result of a variety of estimates, sometimes propagated through chains of publications from early sources (see Supplementary Information). At an order-of-magnitude level (Johnston et al., 2014), one damage event (including single-strand breaks, deaminated cytosines, depurinations, 8-oxo-dGs and other lesions) per mtDNA per day is consistent with several of these arguments, as described in Supplementary Information. We adopt this damage rate by default for our model mtDNA under BER and will allow it to vary in different instances of our simulations. For the model under template repair, we use an order-of-magnitude lower initial estimate of 0.1 damage events per mtDNA per day, reflecting the fact that this pathway is hypothesized to address many replication-induced errors (including mismatches and small indels) that may occur on a slower (and tissue-dependent) timescale relative to some forms of oxidative damage (e.g., 8- oxo-dG) that are addressed by BER (see Supplementary Information).

The BER pathway in plants is incompletely characterised (Boesch et al., 2009; Córdoba-Cañero et al., 2010), so we shape our model of the BER pathway around that reported in (Ferrando et al., 2019; Jeppesen et al., 2011). This pathway involves a collection of five steps catalysed by protein subunits, the least numerous subunits of which are often present at low copy numbers in plants (Supplementary Information) around 4 per mitochondrion (measured in dark-cultured cell suspensions in (Fuchs et al., 2020)). A key player in TR, MSH1, is present at a copy number around 2 per mitochondrion and likely acts in dimer form (Fuchs et al., 2020). We set the production and import rates *λ* and *ρ* per unit time in our model to produce these average per-mitochondrion copy numbers.

### Simulation parameters

We use a time unit of 1 h and report results after 1000 time units in our simulations, having found that system behaviour is stabilised at this time (Supp. Fig. 1). We simulate 30 instances of each parameterisation to characterise variability in the behaviour.

### Code availability

All code is freely available at https://github.com/StochasticBiology/plant-mito-sharing. The simulation code is written in & and the visualisation and analysis code is written in R (R Core Team, 2022); libraries *ggplot2* (Wickham, 2016), *ggpubr* (Kassambara, 2020), *ggbeeswarm* (Clarke et al., 2016) and *viridis* (Garnier et al., 2021) are used for visualisation and *dplyr* (Wickham et al., 2023) is used for data curation.

## Results

### Results from probability theory

In the absence of any interactions between biomolecules, the distribution of copy number across mitochondria would be expected to be described by a Poisson distribution: *Po*(*k*; *θ*) = *θ^k^e*^−*θ*^/*k*! for the probability that a mitochondrion contains *k* molecules, given that the average copy number is *θ* per mitochondrion. For an average of 0.33 mtDNA molecules per mitochondrion (Fuchs et al., 2020), the Poisson picture predicts that 71.7% of mitochondria will be empty of DNA, 23.9% will contain a single mtDNA, 3.98% will contain two mtDNAs, and 0.48% will contain three or more. For an average of 2 copies of a protein per mitochondrion (for example, MSH1 (Fuchs et al., 2020), 13.5% of mitochondria would be expected to contain zero of those proteins, 27% to contain one, and 59.4% to contain two or more (and thus to be capable of forming a functional dimer).

For our model TR process, we require an mtDNA to be in a mitochondrion that contains two or more mtDNAs and two or more copies of MSH1 (see Methods). Consider a given mtDNA molecule. The probability that there is at least one more mtDNA in the same organelle is then 1 − *e*^−*θ_DNA_*^, and the probability that there is at least one more mtDNA and at least two copies of a protein is

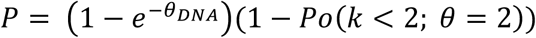

Which is 16.8%, with the low number dominated by the limited number of mitochondria containing two or more mtDNAs. Note that this 16.8% is the percentage of *mtDNA molecules* which can undergo template repair, not the percentage of *mitochondria* with a particular property.

For our model BER process, we require an mtDNA to be in the same mitochondrion as one copy of each of *n* subunits. The Poisson probability of having at least one of each type of protein in a mitochondrion is (1 − *Po*(*k* < 1; *θ*))*^n^*. For a working example, consider a pathway requiring *n* = 5 steps, all involving proteins present with average copy number 4 per mitochondrion. In this case the probability of an mtDNA molecule being colocalised with at least one copy of each of the 5 proteins required is 91%.

The predictions from this simple probability model are illustrated for various different cases in Fig. 2. Clearly, with limited copy numbers of the molecular actors involved, we cannot expect random assignment of biomolecules to mitochondria to support all mtDNA molecules adopting either single-DNA repair mechanisms (like BER) or multiple-DNA, recombination-based repair mechanisms (like TR).

**Figure 2.**
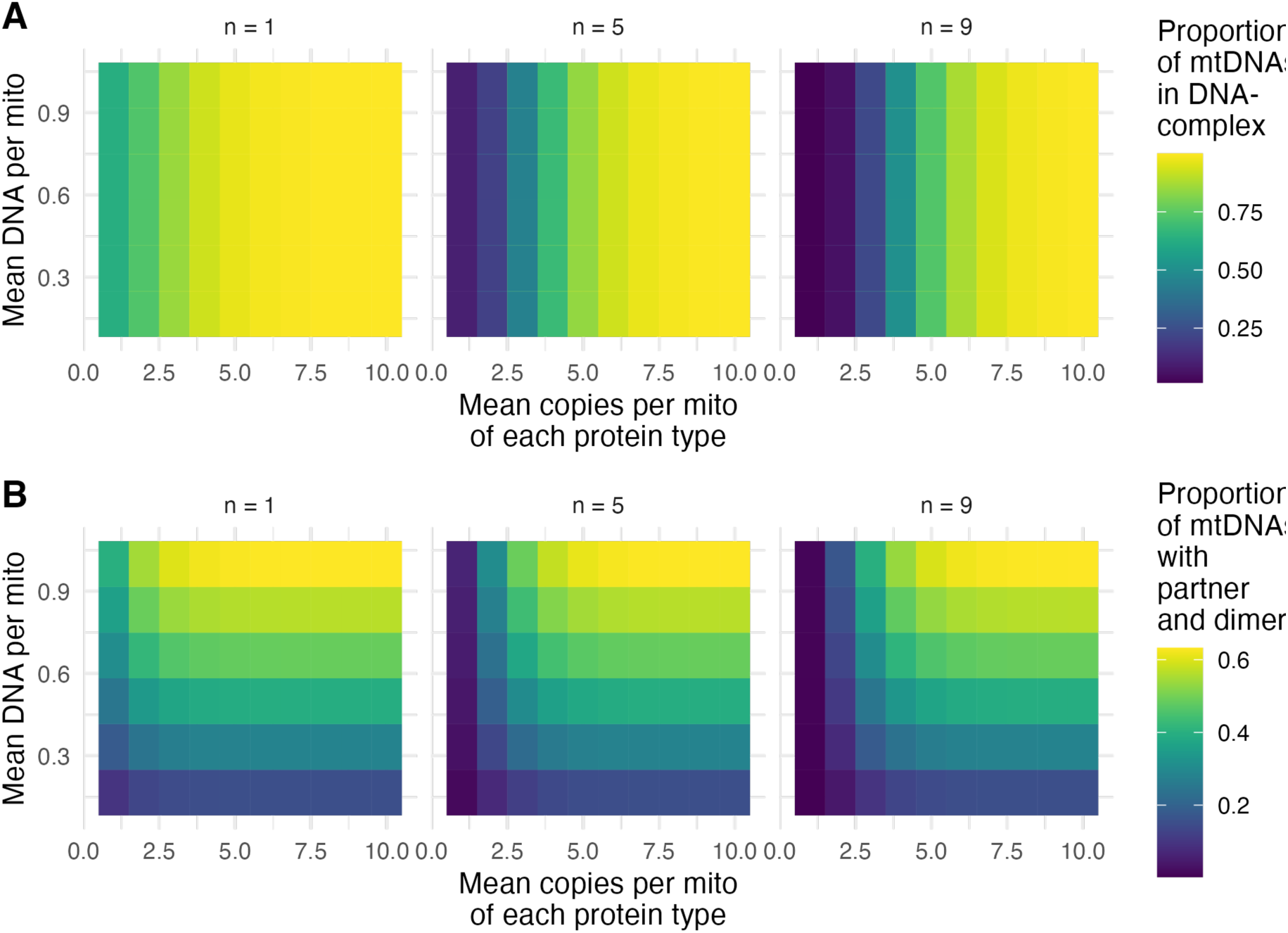
Mathematical results for a simple Poisson model. **(A)** BER-like pathway DNA-complex count and **(B)** TR-like pathway templating capacity for different mean per-mitochondrion occupancies of protein and DNA content. Panels give results for complexes formed of different subunit counts *n*: *n* = 1 describes a single protein being required; higher values describe *n* different subunits required to be present in the same organelle. In (A), repair capacity increases straightforwardly with the probability of a repair complex being present in a mitochondrion. In (B), repair capacity increases with, and is limited by, the number of mtDNAs per mitochondrion, as each mtDNA is required to encounter a partner mtDNA for repair.

### Without fusion-driven exchange or targeted import, DNA repair is compromised

Fig. 3A shows the steady-state proportion of damaged mtDNA molecules in a system without exchange through fusion, where repair is accomplished by the BER pathway. In this case, five steps are involved, each involving a protein present at an average of 4 copies per mitochondrion. With random allocation of proteins to mitochondria, between 2-10% of mtDNA molecules are damaged at a given time. This corresponds roughly to the above 91% probability of an mtDNA being localised with at least one protein for each pathway step.

**Figure 3.**
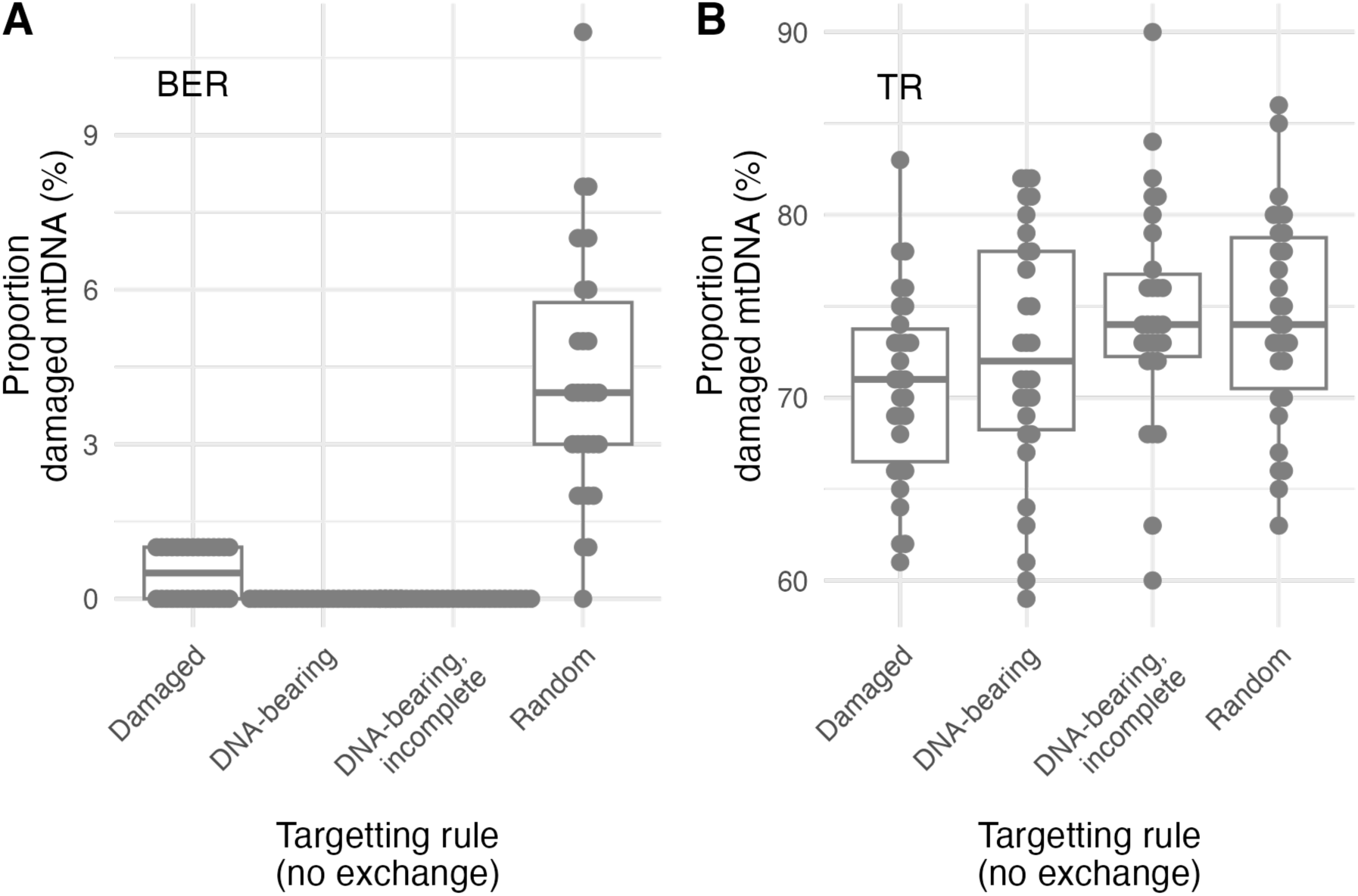
Impact of targetting rules on proportion of damaged mtDNA. Proportion of damaged mtDNA under model **(A)** BER-like pathway and **(B)** TR-like pathway, under different targetting rules while fusion-based exchange is absent. There are 100 mtDNAs in total (so the y-axis can be read as a percentage), and an average 0.33 per mitochondrion.

Fig. 3B shows the same result when repair is accomplished by the TR pathway. Now, in the absence of exchange through fusion, 70-90% of mtDNA is in a damaged state, corresponding roughly to the 16.8% probability above of an mtDNA being colocalised with another mtDNA and a protein dimer.

Targetting protein import specifically to DNA-bearing mitochondria generally decreases the proportion of damaged mtDNA. For the BER pathway this decrease is to almost zero levels; for the template-repair pathway the decrease is much more modest (a few percentage points). In the TR case, the limiting factor is the low probability with which two mtDNAs are present in the same mitochondrion, so protein targetting only has a limited effect. Targetting protein import to mitochondria that have damaged mtDNA is not the best strategy, because only a small subset of mitochondria will contain damaged mtDNA at any given time, and other DNA-bearing mitochondria are then less likely to immediately contain repair proteins when damage does occur.

The proportion of intact mtDNA in these simulations is directly connected to the number of mtDNAs that are present in the same organelle as the set of biomolecules required for repair (either the set of BER proteins, or the TR protein and a partner mtDNA). These proportions, for different targetting rules, are shown in Supp. Fig. 2.

### Exchanging proteins upon fusion helps maintain DNA-repair partnerships; exchanging DNA supports templating repair

Fig. 4A shows the impact on BER-type DNA repair of different fusion and exchange strategies, when the targetting rule is random. If individual proteins are exchanged during fusion events, the proportion of damaged mtDNA drops to be typically under 2%. Exchange of proteins bound together in a metabolon structure – ensuring that a complete set of repair machinery is often present in the same organelle – decreases this damage proportion still further, to the point where no mtDNA is damaged in most cases. Exchanging mtDNA between fused mitochondria, as well as exchanging protein content, presents no particular improvement over exchanging proteins alone, and restricting fusion only to DNA-bearing mitochondria also has little impact.

**Figure 4.**
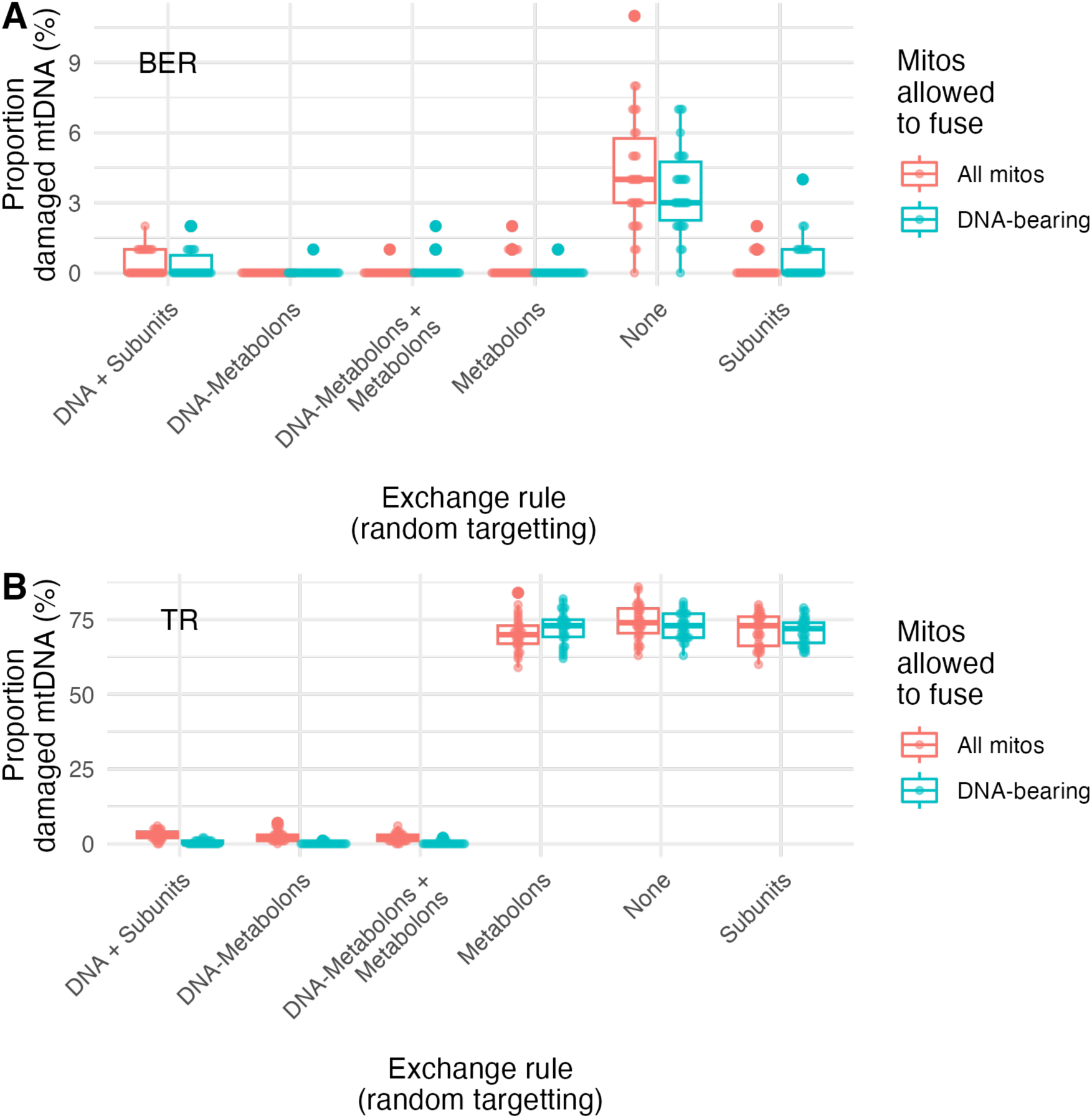
Impact of exchange rules on proportion of damaged mtDNA. Proportion of damaged mtDNA under model **(A)** BER-like pathway and **(B)** TR pathway, under different fusion and exchange rules while targetting is random. In all cases two fusion rules are simulated: allowing fusion between all mitochondria and only allowing fusion between mitochondria containing mtDNA.

The situation with TR (Fig. 4B) is rather different. Now, exchanging proteins provides a slight increase to DNA repair, but the majority of mtDNAs remain unrepaired. However, exchanging mtDNA (with proteins) upon fusion events dramatically improves the repair capacity. If all mitochondria are allowed to fuse, exchanging mtDNA and individual proteins already decreases damage proportion to around 3%; exchanging metabolons (in this case, bound dimers) improves this further to around 2%. The largest improvement is when mtDNA and metabolons are exchanged only between DNA-bearing mitochondria, in which case damage proportion drops to 0.2% or less.

The corresponding statistics for the number of mtDNAs in the same organelle as the set of biomolecules required for repair (either the set of BER proteins, or the TR protein dimer and a partner mtDNA) is shown in Supp. Fig. 3. There is not a direct relationship between the repair capacity and proportion of damaged mtDNA because both statistics reflect snapshots of a dynamic process: an incomplete average repair capacity may still allow effectively all damage to be repaired, depending on the timescales of damage and repair.

### Synergistic contributions of targetting and exchange rules

We next asked if interplay between targetting and exchange (and fusion) rules influenced the proportion of damaged mtDNA in the system (Fig. 5). The model did not demonstrate particularly strong interaction effects between targetting and exchange strategies: beneficial targetting or exchange choices perform well for the BER-like pathway, while only exchange rules have notable influence on the efficacy of the TR-like pathway. The improvement of the TR pathway upon restricting fusion to only mitochondria bearing mtDNA is clear.

**Figure 5.**
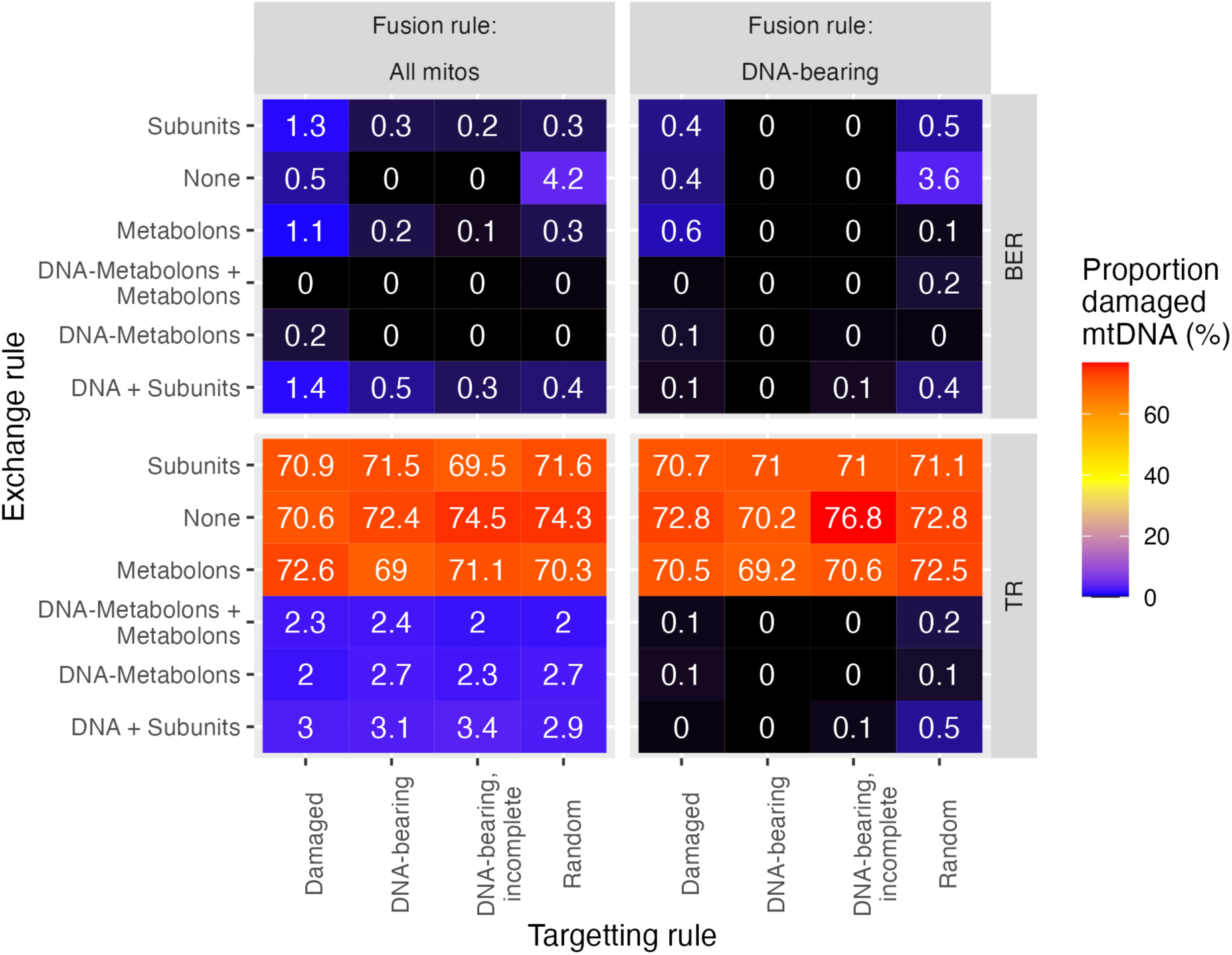
MtDNA damage repair under combined targetting, exchange, and social rules. Each point gives the average percentage of damaged mtDNA in model cells under a given combination of targeting, fusion, and exchange rules.

### Impact of other parameters

Many of the quantities in our model are estimated rather than precisely defined by existing data. Two important quantities which are only roughly estimated are the rate of DNA damage and the “scanning time”, corresponding to the timescale on which damage is recognised by a repair protein colocalised with an mtDNA molecule (Supplementary Information).

In Supp. Fig. 4 we show the effect that varying these parameters has on the proportion of damaged mtDNA in our model. In both cases, the quantitative proportions change in intuitive directions: higher damage rates lead to higher damage proportions, and longer delays in damage recognition and repair also lead to higher damage proportions. However, all values we used for these parameters preserved the qualitative structure of the results in Fig. 5: there were no situations that changed the relative damage rates across the different targetting, social, and exchange rules we considered.

## Discussion

We have shown that the limited copy numbers of repair proteins in individual plant mitochondria can limit the capacity of a single, isolated organelle to repair mtDNA damage. Using estimated damage rates, our model suggests that exchange of repair machinery via fusion and fission between mitochondria in the cellular population is important to maximise mtDNA repair capacity. The physical arrangement of mitochondria shapes the relative weighting of different repair pathways in the cell. Specifically, model BER can keep ∼95% of mtDNA intact without particular targetting or exchange rules; either targetting protein import to DNA-bearing mitos or exchanging protein content drives this to effectively 100%. Model TR requires exchange of contents between mitochondria, largely limited by the colocalisation of two mtDNA molecules in the same organelle. With exchange, especially between only mtDNA-bearing mitochondria, TR can also achieve 100% intact mtDNA. Exchanging of metabolons (collections of repair protein complexes) increases repair efficiency across all circumstances.

The specific proportions of intact mtDNA in our model are dependent on our choices of model parameters, which in some cases are imprecise (including damage rates and protein turnover, described in Supplementary Information). However, the trends and directions of effects in our model are robust to these values: targetted import and exchanging protein content will always drive BER efficiency higher, and exchange – particularly between DNA-bearing mitochondria – will always drive TR efficiency higher regardless of the specific damage rates involved.

Given our observed dependence on the physical arrangement of mitochondria, we can attempt to connect these findings with other research on plant mtDNA. A developing body of theory and experiment suggests that recombination-mediated repair may be particularly important in the shoot apical meristem (SAM) for maintaining the integrity of germline mtDNA (Broz et al., 2022, 2024; Chustecki & Johnston, 2024; Edwards et al., 2021; Johnston, 2019). Indeed, MSH1 – facilitating recombination-mediated repair – seems to be required for germline mutation rate suppression in plants (Broz et al., 2025; Wu et al., 2020). Templated repair is fundamentally limited by the number of mtDNAs in the same compartment as MSH1 dimers. In the SAM, MSH1 expression is high and large-scale fusion brings mtDNAs together in the same compartment (Edwards et al., 2021; Segui-Simarro et al., 2008; Zwonitzer et al., 2024). This recombination can both repair and segregate damage but comes at the cost of structural rearrangements; in somatic tissue, mitochondria remain more fragmented (Arimura, 2018; Chustecki et al., 2021; Logan, 2010) and BER-like pathways, without the risk of structural rearrangements, may be favoured. Pursuing this physical-genetic link, mtDNA-bearing mitochondria have been observed to be more motile and participate more in the “social network” of biomolecular exchange in the mitochondrial population (Chustecki et al., 2025).

We have assumed a “well-mixed” picture inside each mitochondrion. In reality, internal structure limits the mobility of mitochondrial contents, especially mtDNA molecules. Results from yeast emphasise the role of cristae structure in enforcing a “local region” for an mtDNA molecule (Jakubke et al., 2021). There is substantial evidence for recombination-mediated repair in plant mtDNA (Broz et al., 2022, 2024; Gualberto & Newton, 2017), implying an ability to physically colocalise. Recent theoretical work has explored the transport of diffusible contents in fragmented and fused mitochondria (Chuphal et al., 2024; Holt et al., 2024, 2026; Scott et al., 2021). However, whether physical barriers to the colocalisation and movement of plant mtDNA – which may be tethered to the inner membrane and otherwise constrained – exist and impact the rates of our model processes must be the target for future experimental work.

We have not considered the relationship between mtDNA replication and damage/repair dynamics. It is possible that mtDNA replication rates are heterogeneous throughout the cell – for example, they may be elevated in the large-scale network in the SAM. In cases where replication is spatially biased towards regions of efficient repair (like the network), the cell may avoid the accumulation of DNA damage by preferentially replicating mtDNAs that have access to repair machinery.

The maintenance (or abandonment) of mtDNA in plant cells is a much-discussed topic (Oldenburg & Bendich, 2015), and our model cannot substitute for direct experimental investigation of the ideas involved. To this end, new plant lines with diverse organelle mutational profiles will help test the ideas developed here. In particular, advances in technologies that can introduce DNA damage in a targeted fashion (Zhou et al., 2026) have the potential to disentangle the relative prevalence and repairability of mutational damage in germline versus somatic tissues. For now, we hope that this theoretical work helps shed quantitative light on the challenges and possible resolutions involved in plant mtDNA maintenance.

## Acknowledgments

This project has received funding from the European Research Council (ERC) under the European Union’s Horizon 2020 Research and Innovation Programme [Grant agreement No. 805046 (EvoConBiO) to I.G.J.]. This work was supported by the Research Council of Norway through the FRIPRO program [project numbers 357812 and 360449 (MitoPhyto) to I.G.J.] and by the US National Institutes of Health [R35GM148134 to D.B.S.].

## Supplementary Information

### Estimating rates of (mt)DNA damage and repair

Formation of apurinic/apyrimidinic sites, which often convert to single strand breaks, have been measured at around 1.5 sites per 10^6^ bases per day under (human) physiological conditions (Nakamura et al., 1998).

(Saul & Ames, 1986) find: “Based upon measurements of DNA oxidation products in human urine, we estimate that the total rate of formation of all oxidative DNA damage products… is on the order of a few thousand events per cell per day.” This (given around 3 ×10^9^ bases of DNA in the human cell) gives roughly the same scale of estimate of 1 damaged product per 10^6^ bases per day.

(Crine & Verly, 1976) observed that T7 phage DNA in a model of intracellular medium experienced 0.018 SSBs per hour. This was (given about 40 kb in T7 phage DNA) cited with a characteristic SSB timescale of 2.5 h per 10^6^ dsDNA bases in (Shapiro, 1981) – in a table summarising evidence of other modes of damage, of which SSBs are the most common. This figure – roughly 10 damage events per 10^6^ bases per day -- was taken forward as 55 200 SSBs per human nuclear DNA per cell per day in the widely cited (Tice & Setlow, 1985).

Which is the more appropriate class of figure for plant mtDNA? mtDNA is less packaged and protected than nuclear DNA, but likely more so than viral DNA. In many animals, mtDNA accumulates mutations several-fold more rapidly than nuclear DNA (although much of this might be due to differences in repair rates and not solely differences in damage rates (Broz et al., 2025)). We might therefore expect a figure towards the higher end of this range. For BER, we by default adopt a rate of 1 damage event per mtDNA per day – as the plant mtDNA genome is typically several hundred kb (though individual mtDNA molecules may be smaller), this corresponds roughly to the 10 events per 10^6^ bases per day.

For template repair, we consider the rate at which mismatches form due to mtDNA replication. Eukaryotic DNA replication typically involves 1 mismatch every 10^4^-10^5^ bases (Kunkel & Erie, 2015), with in vitro measurements of plant mtDNA polymerase showing similar error rates (Ayala-García et al., 2018). Plant mtDNA turnover can be fast, on the scale of 1 day (Dai et al., 2005), though in other tissues this process is likely on a slower order of magnitude (Bendich, 2010). In that case, we might expect around 0.1 events per mtDNA genome per day, with this value being highly dependent on the amount of mtDNA replication (if any) occurring in the tissue.

According to (Ahmadi et al., 2018) and (Ahmadi et al., 2021), the scanning speed of DNA glycosylases and DNA endonuclease can be around 0.64 µm²/s. With base pair spacing around 0.34 nm per bp, 1 µm corresponds to around 3000 base pairs, so the diffusion rate is around 5×10⁶ bp²/s. The characteristic timescale needed to cover L = 300 kb of DNA with this diffusion constant D is given by L^2^ / 2D = 9×10^3^ s, or several hours. Of course, sliding, unbinding, and non-local motion may change these dynamics beyond those expected from simple 1D diffusive search. We explore several values for the timescale of the initial damage recognition based on this general scale.

**Supplementary Figure 1.**
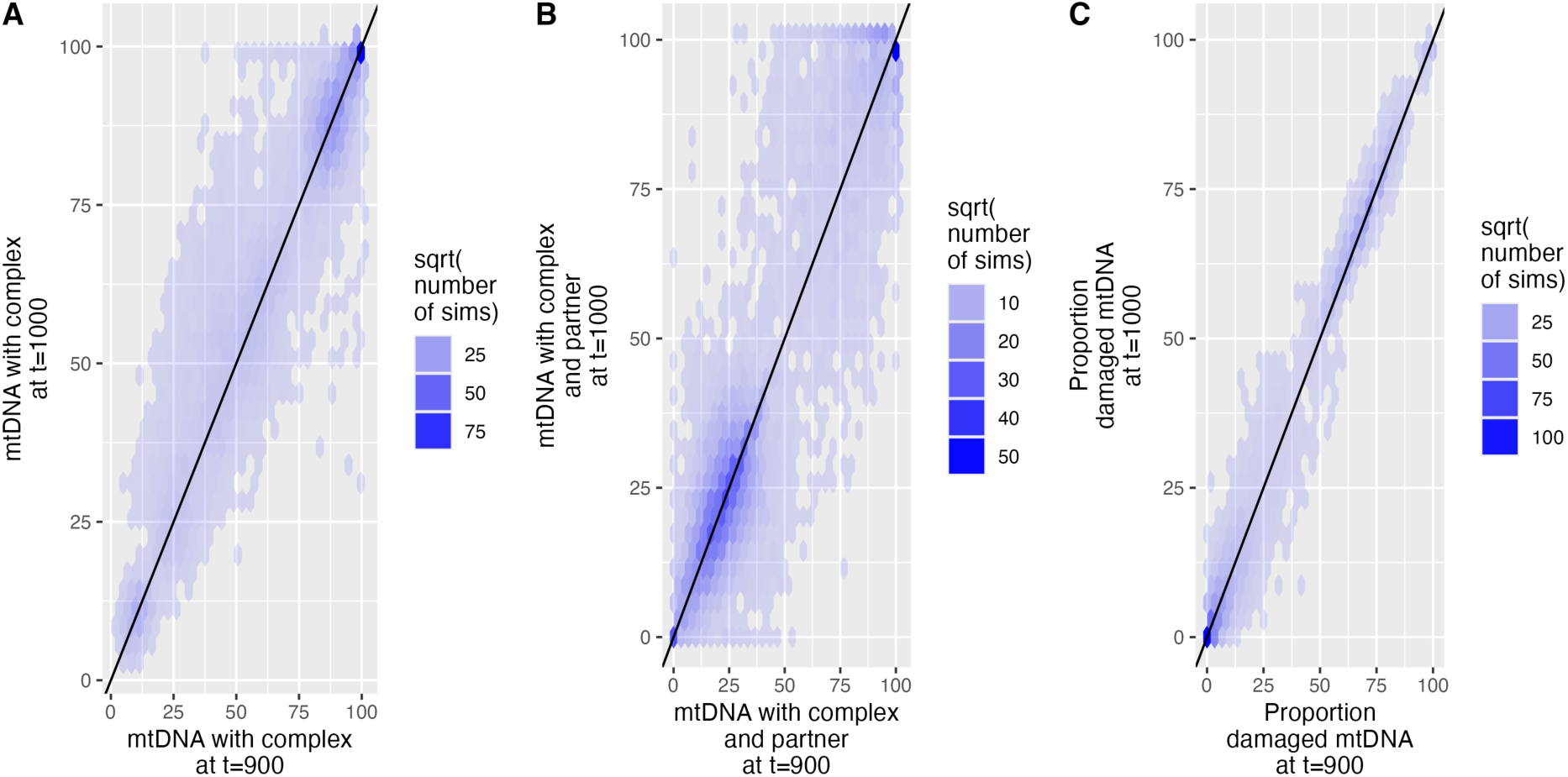
Stability of long-term simulation results. **(A)** DNA-complex count, **(B)** templating capacity, **(C)** proportion of damaged mtDNA at t=900 h and t=1000 h after the start of the simulation. For each quantity, there is strong and symmetric agreement between the earlier and later observations, suggesting stabilisation of the system’s behaviour.

**Supplementary Figure 2.**
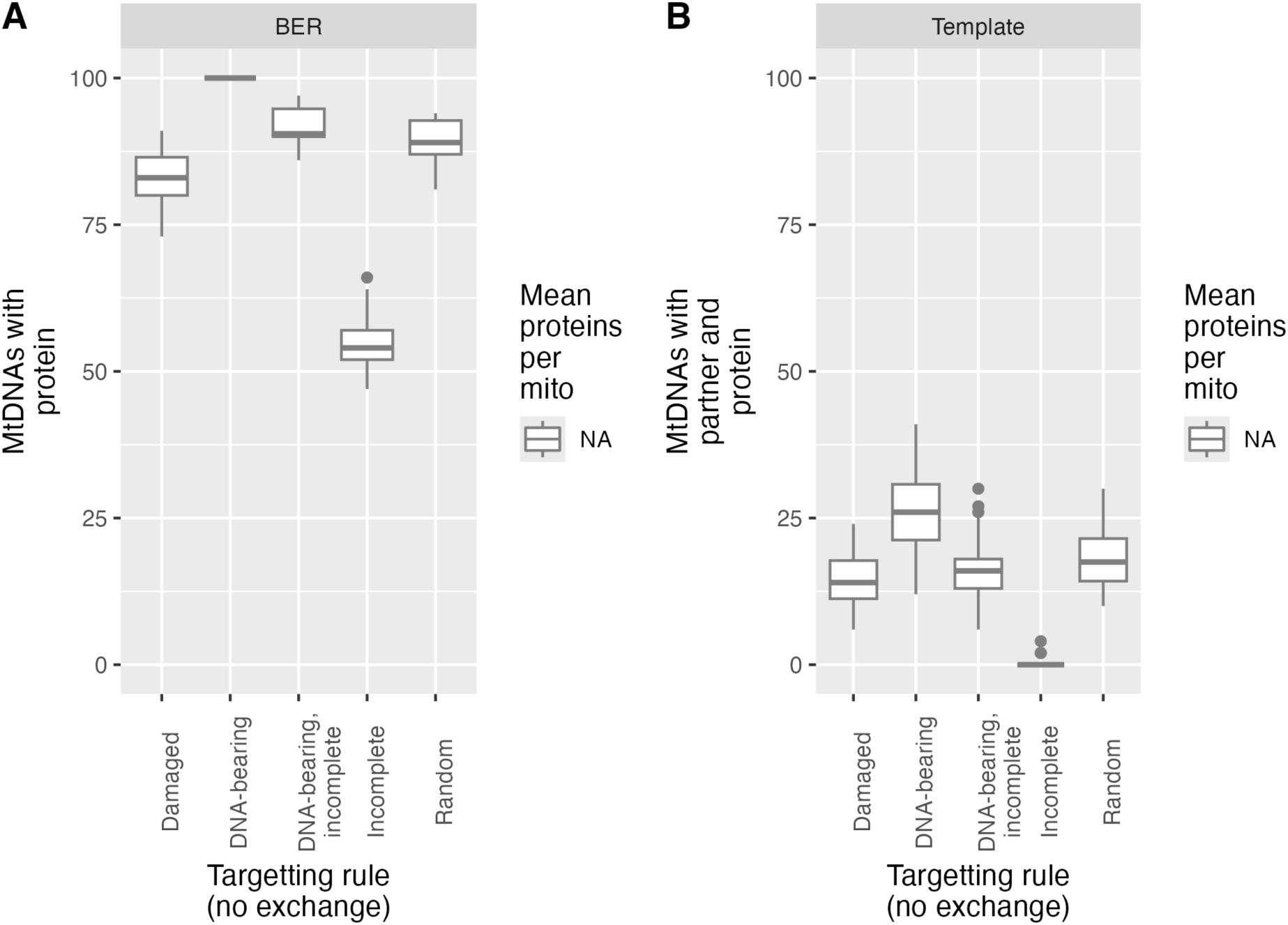
Impact of targetting rules on DNA-complex count and templating capacity. **(A)** DNA-complex count (where a DNA-complex is an mtDNA and one copy of each of the five limiting proteins in the repair pathway) and **(B)** templating capacity (mtDNAs with a dimer of the repair protein and a partner mtDNA), under different targetting rules while fusion-based exchange is absent. There are 100 mtDNAs in total (so the y-axis can be read as a percentage), and an average 0.33 per mitochondrion. Two parameterisations are used, corresponding to an average of 2 and 4 proteins of each type per mitochondrion.

**Supplementary Figure 3.**
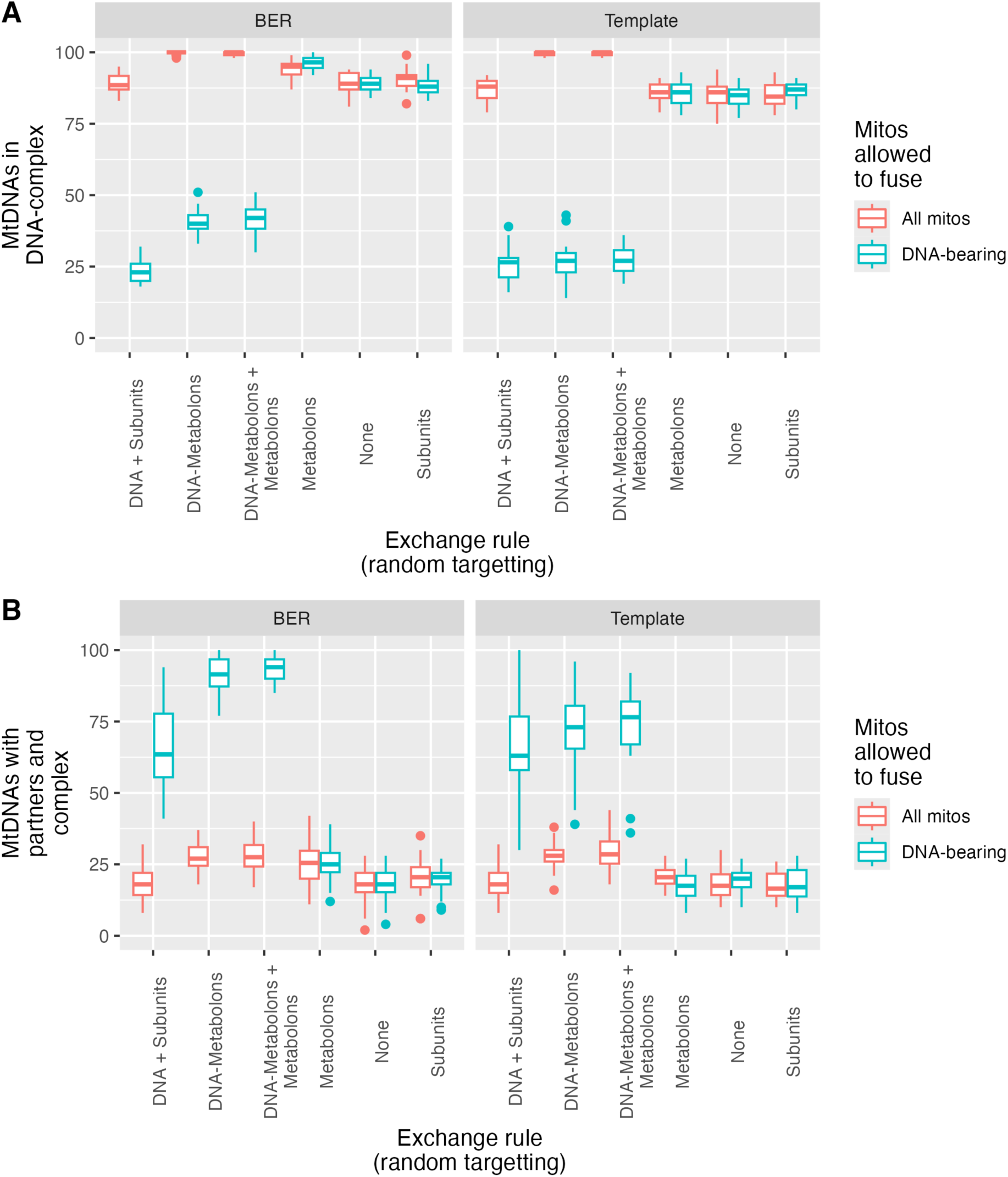
Impact of exchange rules on DNA-complex count and templating capacity. **(A)** DNA-complex count (where a DNA-complex is an mtDNA and one copy of each of the five limiting proteins in the repair pathway) and **(B)** templating capacity (mtDNAs with a dimer of the repair protein and a partner mtDNA), under different exchange rules with random import targetting. There are 100 mtDNAs in total (so the y-axis can be read as a percentage), and an average 0.33 per mitochondrion. Two fusion rules are used: all mitochondria can fuse, or only mitochondria bearing mtDNA.

### Stoichiometries and turnovers of repair proteins

(Ferrando et al., 2019) describes a base-excision repair pathway involving five steps. Of the proteins involved in these five steps that are found in (Fuchs et al., 2020), the minimal copy number element for each step is:

| Step | Protein | Per-mito copy number |
| --- | --- | --- |
| Glycosylase | APS3 Pseudouridine synthase/archaeosine transglycosylase-like family protein | 4.79 |
| Endonuclease/polynucleotide kinase (PNK)/phosphatase | Mitochondrial nuclease 1 (MNU1); Putative endonuclease or glycosyl hydrolase | 4.25 |
| dRP lyase | (none found) |  |
| DNA polymerase | Organellar DNA polymerase gamma I (POLGAMMA1) | 2.48 |
| DNA ligase | DNA ligase 1 (LIG1) | 42.0 |

MSH1 is a key player in plant mtDNA recombination and is likely to play a central role in templated repair (Broz et al., 2022, 2024; Wu et al., 2020; Zwonitzer et al., 2024). (Fuchs et al., 2020) find only two MSH1 copies per mitochondrion. The protein likely acts in dimeric form (Fuchs et al., 2020; Sloan et al., 2025) (Supp. Fig. 5).

(Nelson et al., 2013) looked across proteins and found, with substantial heterogeneity, an average detected protein turnover rate around ∼ 0.1 per day.

**Supplementary Figure 4.**
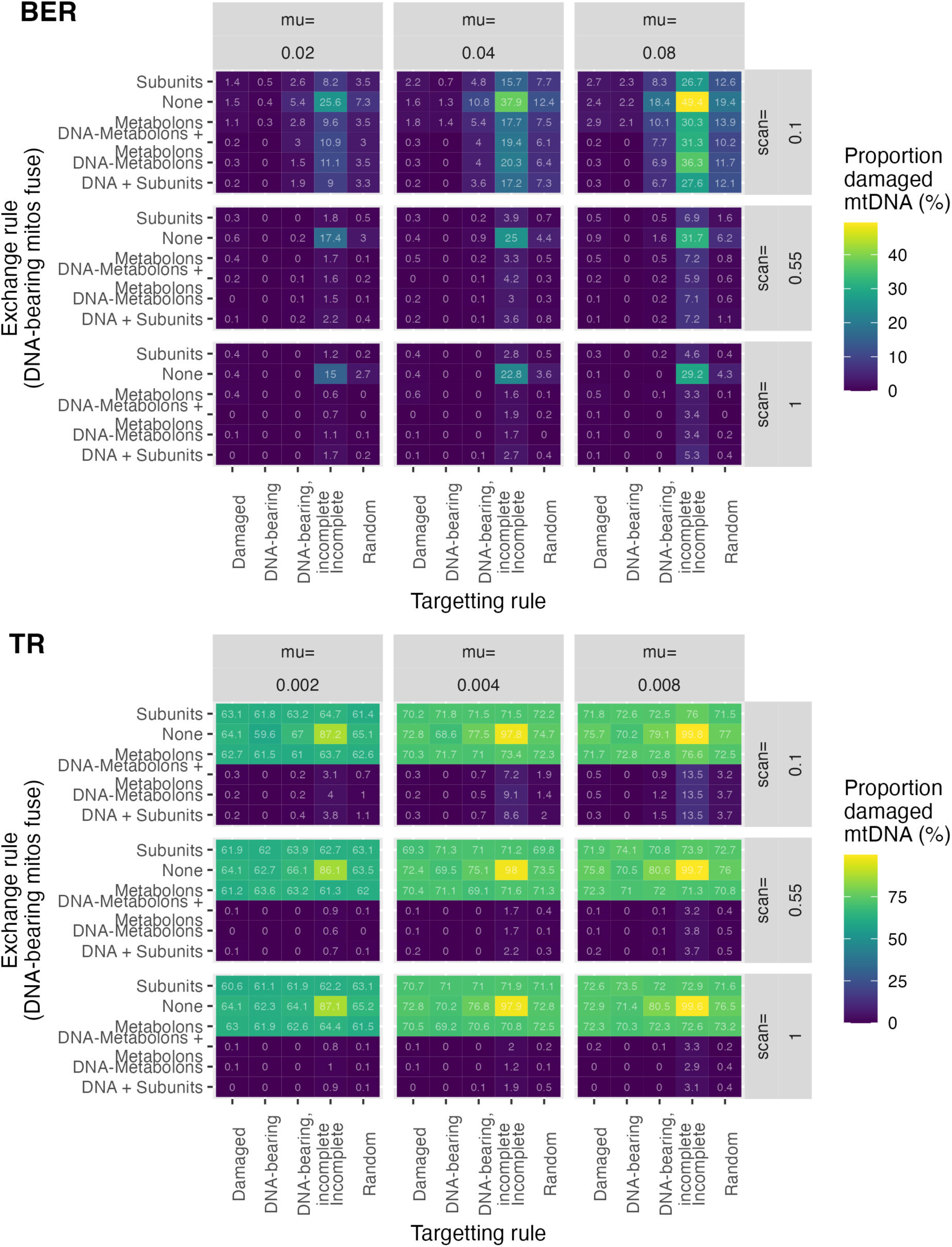
Influence of damage rate (“mu”, per hour) and scanning rate (“scan”, per hour) for (top) BER pathway; (bottom) template repair pathway. Model damage rates are an order of magnitude lower in the TR experiments (see text). Each panel gives a combination of these parameters; each pixel in a panel (as in Fig. 5) gives the proportion of damaged mtDNA under those conditions.

**Supplementary Figure 5.**
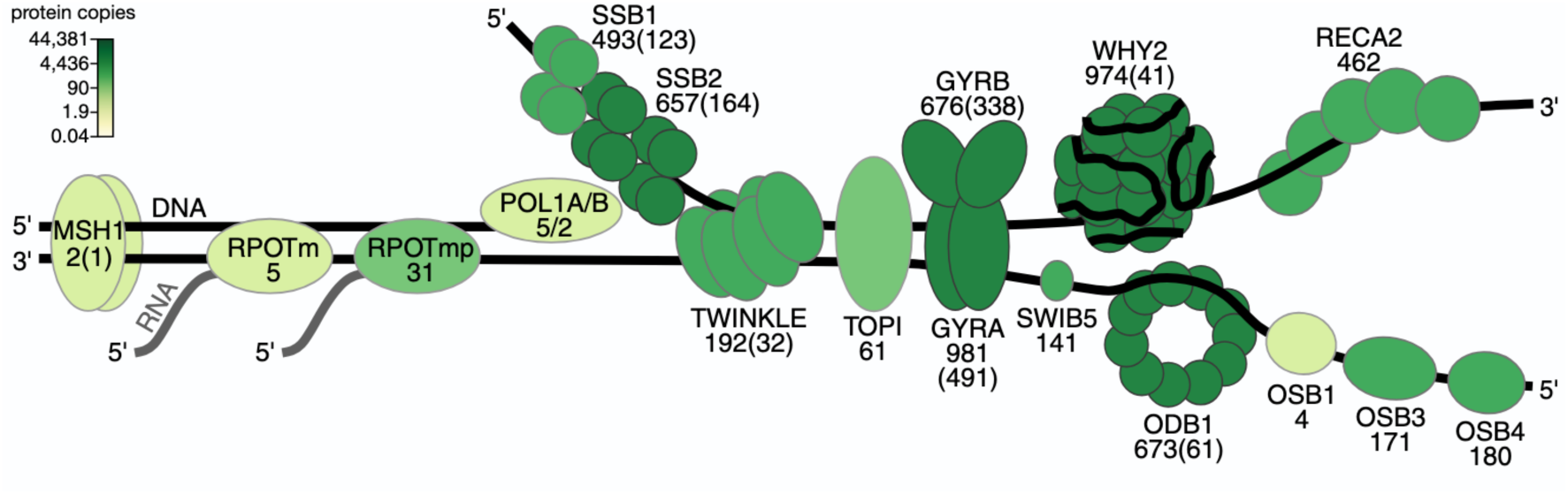
Per-mitochondrion copy numbers of mtDNA-associated protein from (Fuchs et al., 2020) (under CC-BY).

## Notes

https://github.com/StochasticBiology/plant-mito-sharing

### Competing Interest Statement

The authors have declared no competing interest.

